# 16S rRNA gene sequencing for bacterial identification and infectious disease diagnosis

**DOI:** 10.1101/2024.10.14.618149

**Authors:** Mei-Na Li, Qiang Han, Nan Wang, Ting Wang, Xue-Ming You, Shuai Zhang, Cui-Cui Zhang, Yong-Qiang Shi, Pei-Zhuang Qiao, Cheng-Lian Man, Teng Feng, Yue-Yue Li, Zhuang Zhu, Ke-Ji Quan, Teng-Lin Xu, George Fei Zhang

## Abstract

16S rRNA gene sequence is the most common housekeeping genetic marker to study bacterial phylogeny and taxonomy. Therefore, 16S rRNA gene sequencing has the potential to identify novel bacteria and diagnose bacteria. This study compared 16S rRNA gene sequencing with conventional PCR for bacterial identification and disease diagnosis. The bacterial community in healthy and diseased hosts was analyzed by 16S rRNA gene sequencing. 16S rRNA gene sequencing is more sensitive than conventional PCR in detecting bacteria. Moreover, 16S rRNA gene sequencing is adequate to identify novel bacteria. 16S rRNA gene sequencing demonstrated that most pathogenic bacteria persist in diseased or healthy hosts in different abundance. Pathogenic bacteria, such as well-known chicken pathogen *Avibacterium paragallinarum*, *Ornithobacterium rhinotracheale*, and *Gallibacterium anatis*, were identified as indicator species of diseased samples. Alpha diversity analysis showed that the healthy group species is significantly higher than in the diseased groups. Beta diversity analysis also demonstrated differences between healthy and infected groups. The study concluded that 16S rRNA gene sequencing is a more sensitive method for detecting pathogens, and microbiota analysis can distinguish between healthy and diseased samples. Eventually, 16S rRNA gene sequencing has represented the potential in human and animal clinical diagnosis and novel bacterial identification.

## 1 Introduction

Viruses and bacteria are usually identified as pathogens of infectious diseases. Bacteria associated with infectious diseases have been identified in human and animal samples. Disease diagnosis is essential to protect both humans and animals against contagious diseases. Polymerase Chain Reaction (PCR) is a well-known, sensitive, specific, and dependable diagnostic method most widely used in clinical laboratories [1]. Detection of pathogens [2,3] and vaccination [4,5] are valuable strategies for preventing infectious diseases, and research has been conducted on the virulence of different pathogens [6–8]. The metagenomic sequencing approach is a supplementary method to point-of-care diagnosis using PCR-based multiplex assays [9].

16S rRNA gene sequencing is a powerful tool for identifying novel pathogens in patients with suspected bacterial disease, and more recently, this technology has been used in clinical laboratories for the routine identification of bacterial isolates [10]. However, 16S rRNA gene sequencing for bacterial identification has been limited to diagnosing human diseases, not animals. Infectious coryza is a significant respiratory tract disease of poultry, which is primarily caused by *Avibacterium paragallinarum* (APG) and *Ornithobacterium rhinotracheale* (ORT) [11,12]. PCR is widely used for disease diagnosis in poultry. However, 16S rRNA gene sequencing is limited for disease diagnosis in poultry.

In this study, 16S rRNA gene sequencing was conducted for disease diagnosis and bacterial identification. Seven groups (healthy and diseased) were analyzed by 16S rRNA gene sequencing. For disease diagnosis, the result of 16S rRNA sequencing was compared with that of conventional PCR. The bacterial community composition of each group was demonstrated. Indicator species, alpha diversity, and beta diversity were examined to illustrate the differences between health and disease.

## 2 Materials and Methods

### 2.1 Study design and sample collection

In this research, 35 swab samples were obtained from the suborbital sinuses of chickens with swollen head syndrome (SHS) and specific-pathogen-free chickens. These swab samples included five specific-pathogen-free samples (Mock), five samples artificially infected with APG (C-APG), five samples artificially infected with ORT (C-ORT), seven samples testing positive for APG (P-APG), three samples testing positive for ORT (P-ORT), three samples testing positive for both ORT and APG (P-APG-ORT), and seven samples tested negative for APG and ORT (N-APG-ORT). All samples underwent initial testing using conventional PCR. These samples from clinically affected laying hens were examined using PCR to determine the presence of APG and ORT [13]. Serovar A, B, and C of APG were identified by multiple PCR using previously described primers. PCR identified ORT with primers [13]. The Animal Care and Use Committee of Wohua Biotech reviewed and approved the animal experiments. They were conducted under the guidelines for experimental animals of the Ministry of Science and Technology of the People’s Republic of China.

### 2.2 DNA extraction, PCR amplification, and 16S rRNA gene sequencing

Microbial DNA was extracted using the HiPure Universal DNA Kits (Magen, China) following the manufacturer’s protocols. The V3-V4 region of the ribosomal RNA gene was amplified by PCR, as described previously [13]. The amplification used primers 341F and 806R [13]. The amplicons were extracted from 2% agarose gels and purified using the AxyPrep DNA Gel Extraction Kit (Axygen Biosciences) following the manufacturer’s instructions. The quantification was performed using the ABI StepOnePlus Real-Time PCR System (Life Technologies). Subsequently, the purified amplicons were equimolarly pooled and subjected to paired-end sequencing (2X250) on an Illumina platform following standard protocols. The raw reads have been submitted to the NCBI Sequence Read Archive (SRA) database under the Accession Number PRJNA1165975.

### 2.3 Bioinformatic analysis

The clean tags were clustered into operational taxonomic units (OTUs) of ≥ 97% similarity using the UPARSE pipeline. The representative sequences were classified into organisms by a naive Bayesian model with the RDP classifier based on the SILVA database. The abundance statistics for each taxonomy were visualized using Krona. The community composition was represented in a stacked bar plot using the R project’s ggplot2 package. Alpha diversity index (Sobs) comparisons between groups were calculated using Welch’s t-test. Beta diversity analysis was conducted, and sequence alignment was performed using Muscle. A phylogenetic tree was constructed using FastTree. Multivariate statistical techniques, including PCoA (principal coordinates analysis) of Bray-Curtis distances, were generated and plotted using the Vegan package and the ggplot2 package in the R project. Statistical analysis using Welch’s t-test was conducted in the R project’s Vegan package.

### 2.4 Statistical analysis

Statistical analysis and plotting were conducted using GraphPad Prism 9. Statistical analysis between three or more groups was performed using one-way ANOVA with Dunnett’s multiple comparison test. P < 0.05 stands for a significant difference. “ns” indicates not significant, and “*” indicates P < 0.05.

## 3 Results

### 3.1 16S rRNA gene sequencing is efficient in disease diagnosis and novel bacteria identification

16S rRNA gene sequencing is quantifiable and more sensitive than conventional PCR. 16S rRNA gene sequencing demonstrated that APG was detected in the group that tested positive for APG using PCR and in the groups that tested negative for APG using PCR (Figure 1). When compared to the SPF chicken (Mock), a higher relative abundance of APG was observed in P-APG, C-APG, C-ORT, P-ORT, P-APG-ORT, and N-APG-ORT (Figure 1A). Furthermore, ORT was also identified in each group by 16S rRNA gene sequencing (Figure 1A). Clinical Samples with SHS showed upregulated bacterial community composition of *Psychrobacter*, *Acinetobacter*, *Pseudomonas*, and *Escherichia*-*Shigella* genera levels compared to SPF and artificially infected chickens (Figure 1B). The relative abundance of APG in the C-APG, C-ORT, P-APG, P-ORT, P-APG-ORT, and N-APG-ORT groups was higher than in the mock group (Figure 1C). Notably, the relative abundance of APG in the P-APG group was significantly higher than in the mock group (Figure 1A). Additionally, a higher abundance of ORT was observed in C-ORT, P-ORT, P-APG, P-APG-ORT, and N-APG-ORT compared to the Mock or C-APG groups (Figure 1B). In the APG testing, 15 out of 35 samples tested positive using conventional PCR, while all 35 samples tested positive for APG through 16S rRNA gene sequencing (Figure 1E). Similarly, for ORT testing, 11 out of 35 samples tested positive using conventional PCR, and all 35 samples tested positive for ORT using 16S rRNA gene sequencing (Figure 1E).

**Figure 1.**
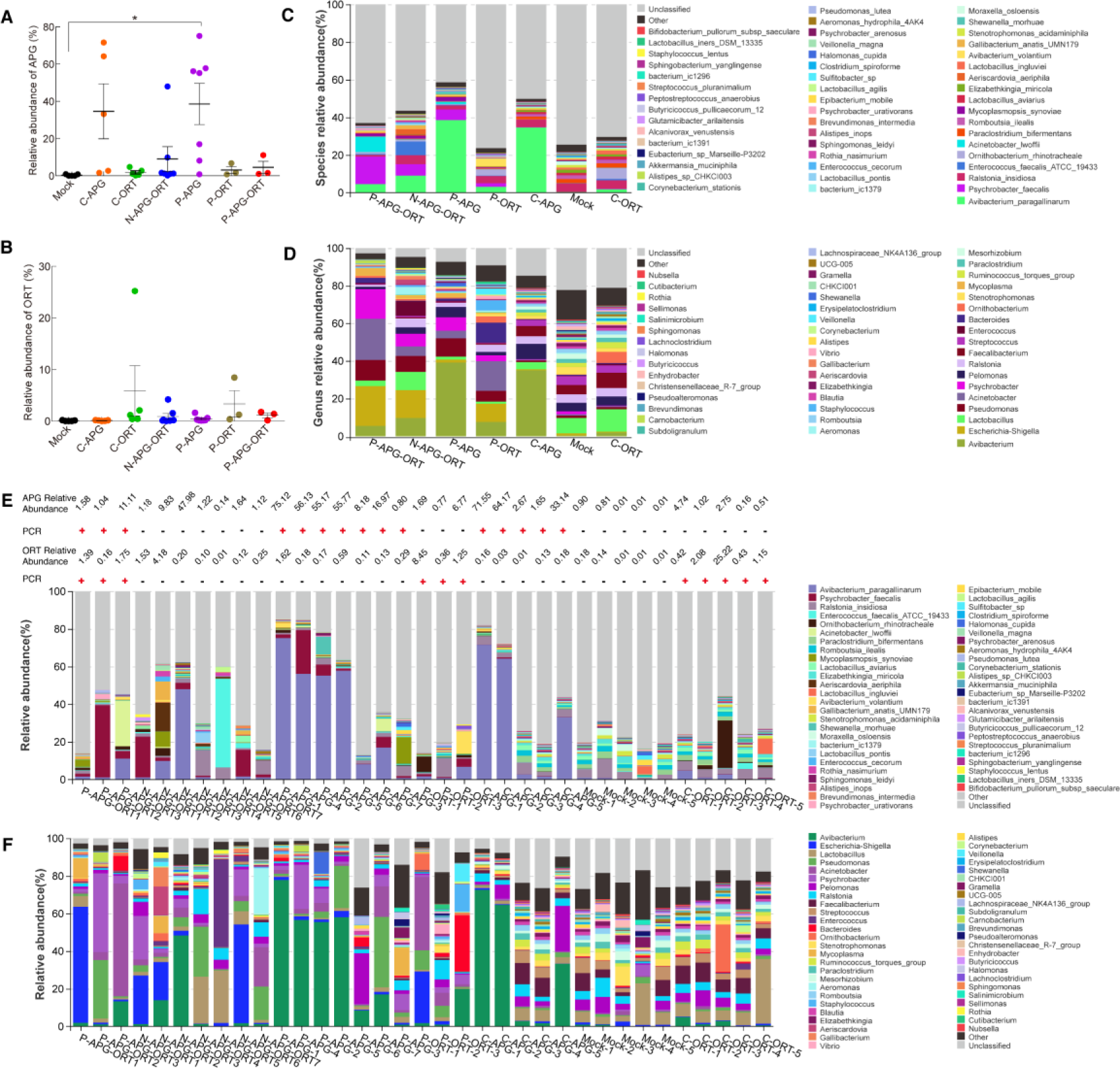
The microbiota of the samples was analyzed using 16S rRNA sequencing. This study included seven groups: Mock (specific pathogen-free), C-APG (artificially infected with APG), C-ORT (samples artificially infected with ORT), P-APG (APG positive samples), P-ORT (ORT positive samples), P-APG-ORT (APG and ORT positive samples), and N-APG-ORT (APG and ORT negative samples). (A) Shows the relative abundance of species in chickens from groups P-APG-ORT, N-APG-ORT, P-APG, P-ORT, C-APG, Mock, and C-ORT. (B) Shows the relative abundance of genera in chickens from the same groups. (C) Depicts APG relative abundance in different groups. (D) Depicts ORT relative abundance in the groups. (E) Presents the 16S rRNA sequencing and conventional PCR data for all the samples.

### 3.2 Composition of bacterial community in health and disease

The study evaluated the community composition of healthy and diseased groups, focusing on the top 50 species and genera. The dominant bacterial community in SPF animal samples was *Ralstonia insidiosa*, *Paraclostridium bifermentans*, *Romboutsia ilealis*, and *Lactobacillus aviaries* (Figure 1A). These bacteria were also predominant in the APG and ORT infection groups, with a higher abundance in the P-APG and C-APG groups. Animals with SHS showed a higher APG abundance and different microbiota than healthy animals (Figure 1). Additionally, *Mycoplasmopsis synoviae*, *Enterococcus faecalis*, *Psychrobacter faecalis*, and *Acinetobacter lwoffii* were detected in nature-infected animals with SHS. At the genus level, *Streptococcus*, *Faecalibacterium*, *Ralstonia*, *Pelomonas*, *Psychrobacter*, *Acinetobacter*, *Pseudomonas*, *Lactobacillus*, and *Escherichia-Shigella* were found to be predominant in healthy animals (Figure 1B). The artificial infection group, C-APG, and C-ORT, shared a similar genus relative abundance with healthy animals but showed an abnormal relative abundance of APG (Figure 1B). Diseased animals with SHS exhibited an upregulated relative abundance of *Psychrobacter*, *Acinetobacter*, *Pseudomonas*, and *Escherichia-Shigella*, and a downregulated relative abundance of *Streptococcus*, *Faecalibacterium*, and *Lactobacillus*.

### 3.3 Indicator species analysis

The Mock group was compared with the diseased group for indicator species identification. Indicator species in infected groups (P-APG, P-ORT, P-APG-ORT, N-APG-ORT, C-APG, C-ORT) or groups with SHS (P-APG, P-ORT, P-APG-ORT, N-APG-ORT) was identified (Figure 2). APG is the indicator species of P-APG, P-APG-ORT, N-APG-ORT, and C-APG groups, while ORT is the indicator species of P-ORT, P-APG, P-APG-ORT, and C-ORT groups. Compared with the Mock group, the indicator species of the diseased groups include *Avibacterium paragallinarum*, *Gallibacterium anatis*, *Ornithobacterium rhinotracheale*, *Avibacterium volantium*, *Peptostreptococcus anaerobius*, *Lactobacillus agilis*, *Veillonella magna*, *Rothia nasimurium*, *Psychrobacter faecalis*, *Aeriscardovia aeriphila*, *Psychrobacter urativorans*, *Psychrobacter arenosus*, *Lactobacillus coleohominis*, *Staphylococcus lentus* (Figure 2).

**Figure 2.**
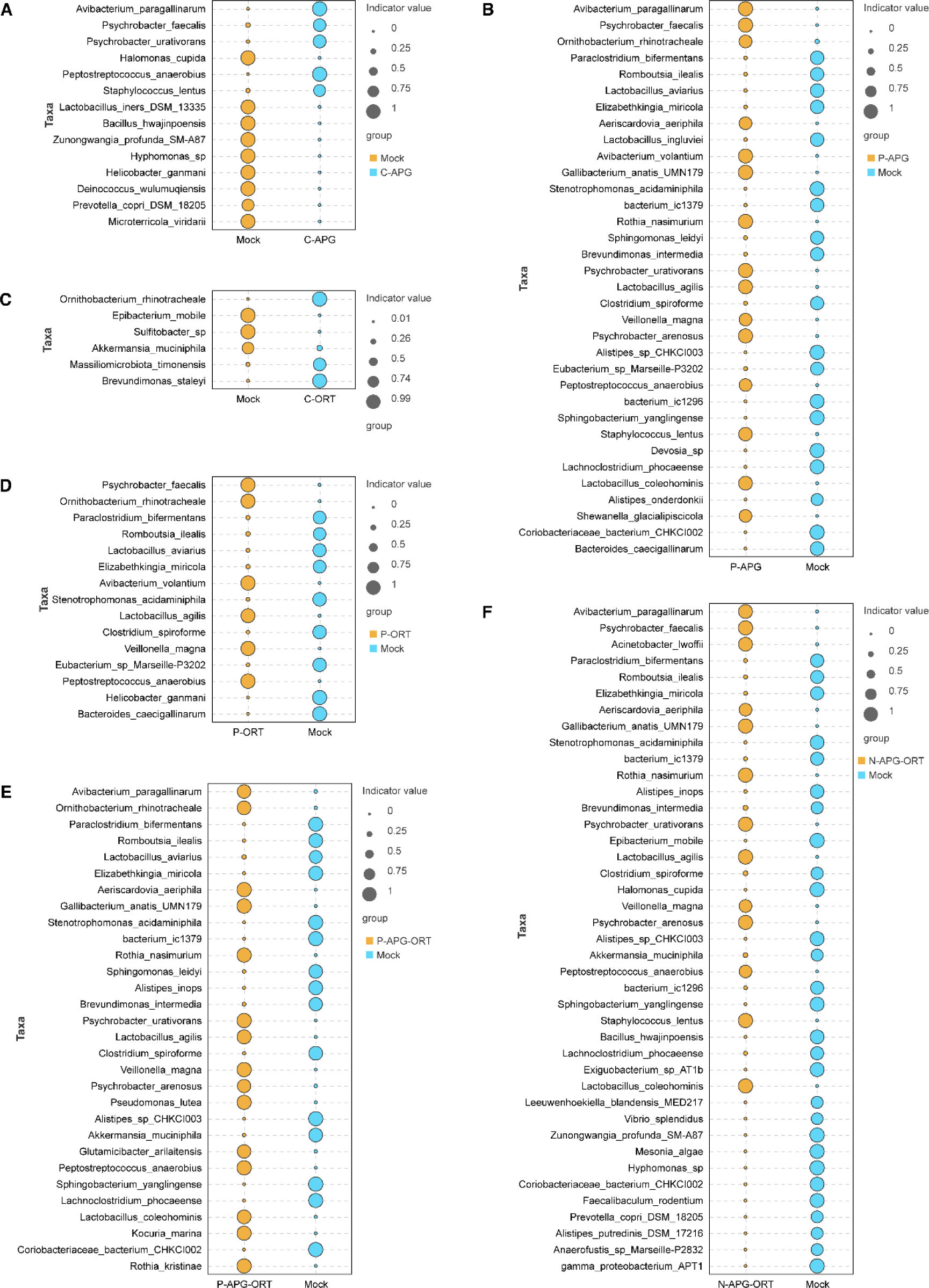
Indicator species analysis. Each diseased group (P-APG, P-ORT, P-APG-ORT, N-APG-ORT, C-APG, C-ORT) was compared with the healthy group (Mock).

Meanwhile, it demonstrated that the indicator species of the Mock group including *Romboutsia ilealis*, *Stenotrophomonas acidaminiphila*, *Elizabethkingia miricola*, *Clostridium spiroforme*, *Romboutsia ilealis*, *Paraclostridium bifermentans*, *Coriobacteriaceae bacterium*, *Lachnoclostridium phocaeense*, *Sphingobacterium yanglingense*, *Alistipes sp CJKCI003*, *Brevundimonas intermedia*, *bacterium ic1379*, *Lactobacillus aviaries*, *Sphingomonas leidyi*, *Alistipes inops*, *Bacillus hwajinpoensis*, *Helicobacter ganmani*, *Akkermansia muciniphila*, *Epibacterium mobile*, *Bacteroides caecigallinarum*, *Eubacterium sp Marseille*, *Halomonas cupida*, and *Lactobacillus iners* (Figure 2).

### 3.4 Alpha diversity analysis

The alpha diversity was assessed by comparing the Sobs index (number of observed OTUs) across different groups. The sobs richness index of all treatments was compared with each other. The sobs richness index of the Mock group was significantly higher than that of the P-APG, P-ORT, C-APG, C-ORT, P-APG-ORT, and N-APG-ORT groups. The sobs richness index of the P-APG group was lower than that of the P-ORT, N-APG-ORT, P-APG-ORT, and Mock groups (p=0.004), as well as the C-APG and C-ORT groups, respectively (Figure 3A-F). Additionally, the sobs richness index of P-ORT was lower than that of P-APG-ORT and the Mock group (Figure 3G-K, ns). Furthermore, the sobs richness index of N-APG-ORT was lower than that of P-APG-ORT, the Mock group, C-APG, and C-ORT (Figure 3L-O). The sobs richness index of P-APG-ORT was lower than that of the Mock group (p=0.0461) and higher than that of C-APG and C-ORT (Figure 3P-R). Finally, the sobs richness index of the Mock group was significantly higher than that of C-APG (p=0.0037) and C-ORT (p=0.0019), respectively (Figure 3S-T). No significant differences were observed between C-APG and C-ORT (Figure 3U).

**Figure 3.**
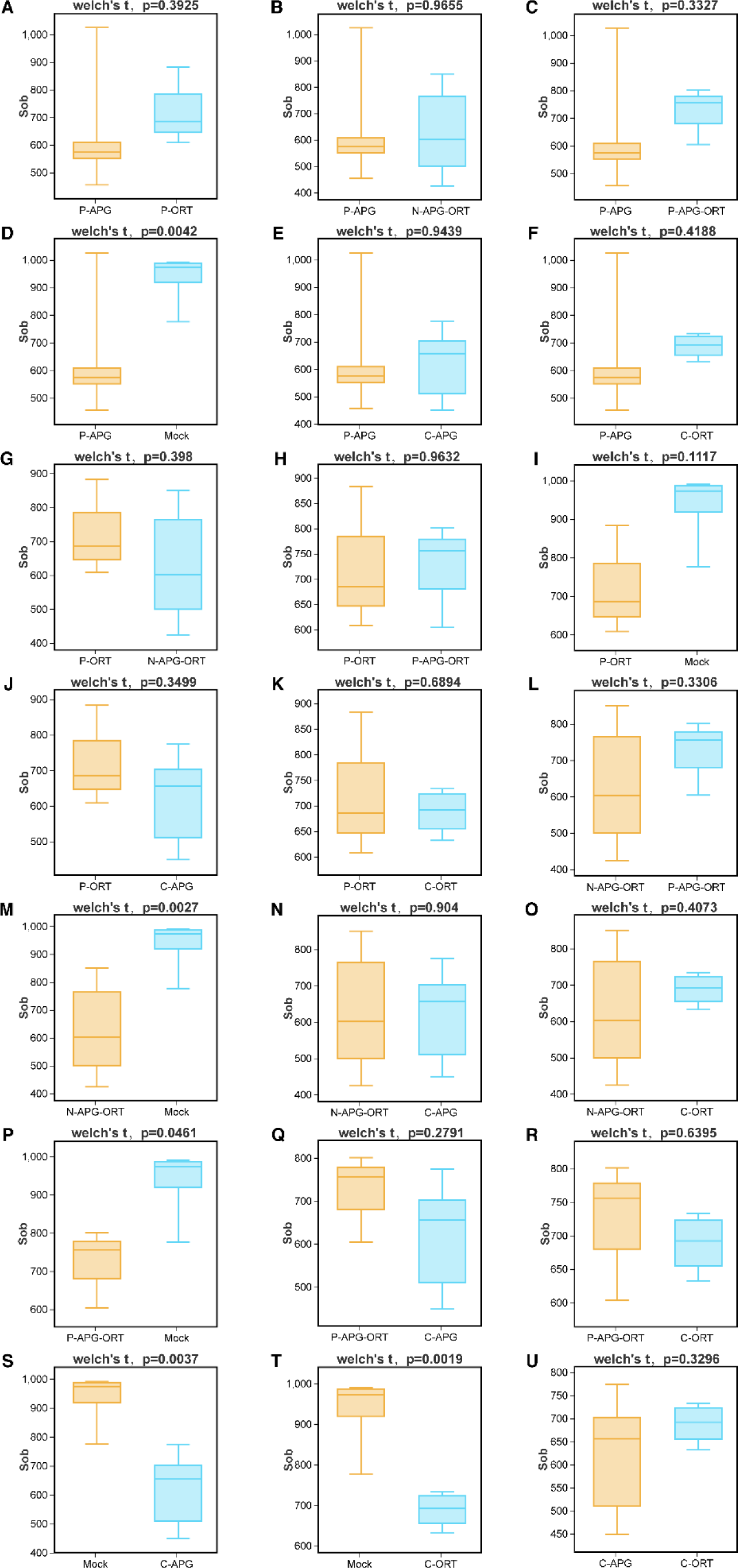
Alpha diversity analysis. The alpha diversity analysis was performed by comparing the Sobs index among different groups. The comparative analysis included Mock, C-APG, C-ORT, P-APG, P-ORT, P-APG-ORT, and N-APG-ORT. Mock’s Sobs index was higher than that of C-APG, C-ORT, P-APG, P-ORT, P-APG-ORT, and N-APG-ORT.

### 3.5 Beta diversity analysis

The beta diversity analysis involved constructing a UPGMA cluster dendrogram and performing PCoA analysis. In the C-ORT group, all the samples were clustered together, as were the samples in the Mock group (Figure 4A). In contrast, the five samples in the C-APG group were spread across two large branches. Similarly, five out of seven samples in the P-APG group were grouped together, while the remaining two were in separate branches. The samples of P-ORT were distributed across three branches, and the N-APG-ORT samples were in different branches. Figure 4B illustrates that the bacterial community structure of C-ORT was most similar to that of the Mock group, although there were differences within the Mock group. The considerable distance among samples in the C-APG, P-APG, P-ORT, N-APG-ORT, and P-APG-ORT groups indicated low similarity within each group (Figure 4B). On the other hand, the proximity of samples in the Mock and C-ORT groups indicated high similarity within each group (Figure 4B).

**Figure 4.**
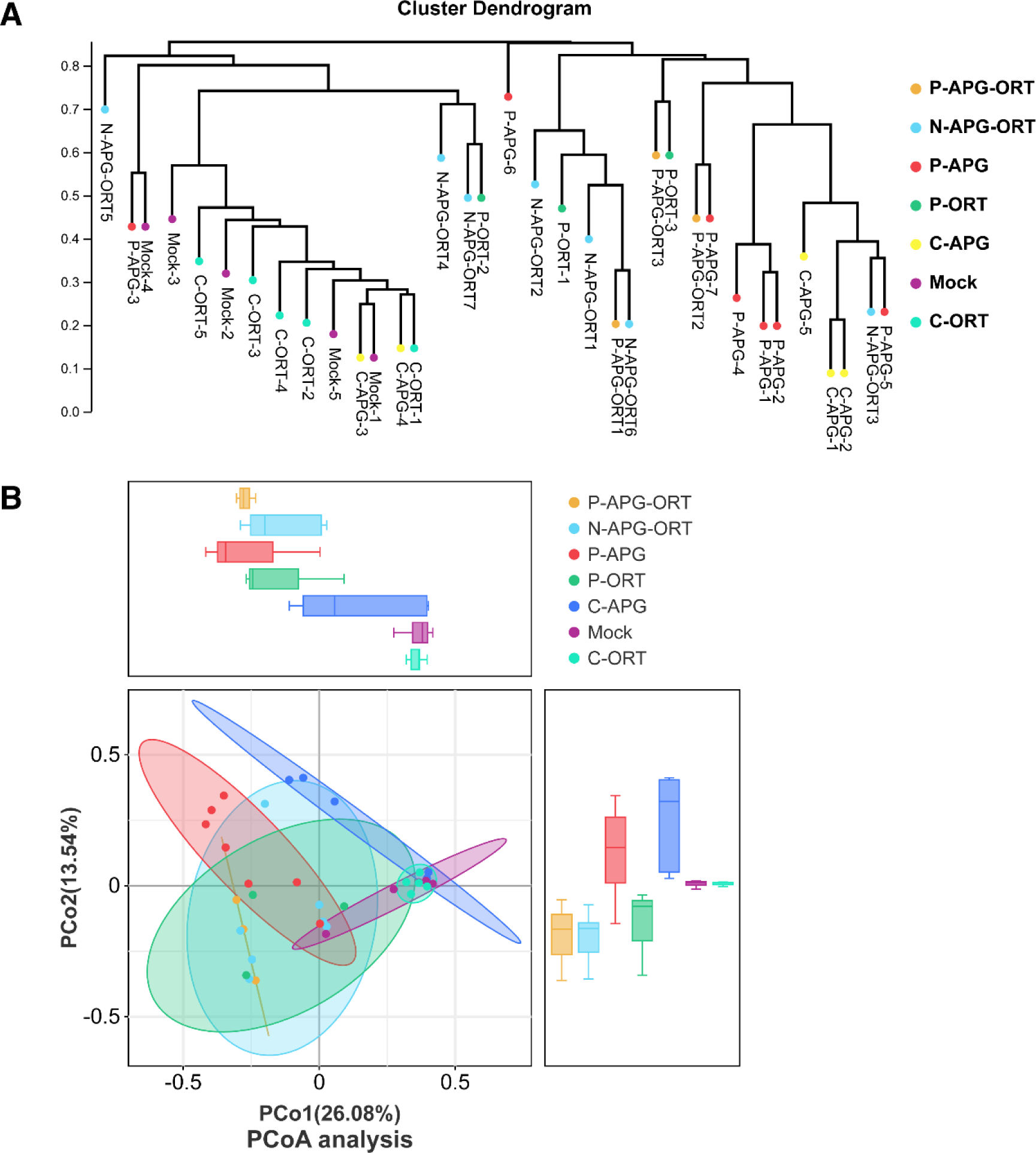
Beta Diversity analysis. Mock group samples are clustered closely on the same branches or nearby. Meanwhile, samples in the C-APG, P-APG, P-ORT, P-APG-ORT, and N-APG-ORT groups are situated on different branches (A). Additionally, samples in the diseased group exhibited a greater separation than those in the healthy group (B).

## 4. Discussion

This research utilized 16S rRNA gene sequencing to identify bacteria in infected animals. 16S rRNA gene sequencing was compared with conventional PCR testing. Conventional PCR has limitations in terms of low sensitivity and often yields false negative results. Bacterial community composition of chickens was analyzed in this study. It’s reported that 16S rRNA is more sensitive and quantifiable than conventional PCR in human clinical diagnosis [14]. Additionally, the study by Fida Madiha and colleagues suggests that 16S rRNA PCR/sequencing is clinically valuable and essential in diagnosing patients with culture-negative infections [15]. This study showed that many bacterial species had significantly higher relative abundances in the SHS groups than in the Mock group. Therefore, conventional PCR is not reliable for clinical diagnosis.

16S rRNA gene sequencing can identify both known and novel bacteria. APG and ORT were identified through this method, along with various normal microbiota and newly discovered pathogenic bacteria. A high similarity of the composition of each sample among Mock groups was determined. Whereas, infected groups showed an entirely different bacterial community composition. The relative abundances of specific species and genera were higher in the diseased groups than in the Mock group. For instance, the relative abundances of *Avibacterium*, *Escherichia-Shigella*, *Pseudomonas*, *Acinetobacter*, and *Psychrobacter* genera were higher in the P-APG, P-ORT, P-APG-ORT, and N-APG-ORT groups than in the Mock group. This study also identified microbiota with higher relative abundances in the P-APG, P-ORT, P-APG-ORT, N-APG-ORT, C-APG, and C-ORT groups than in the Mock group. Previous studies have reported *Staphylococci*, *Streptococci*, and *Enterococci* as regular bacterial species colonizing the trachea [16]. Also, as reported by Shabbir et al., lung microbiota is quite different from that of the trachea and may contain *Avibacterium*, *Pseudomonas*, or *Bordetella* [17]. In general, lung microbiota usually contains *Lactobacillus*, *Enterococcus*, *Streptococcus*, *Staphylococcus*, *Corynebacterium*, *C. disporicum*, *E. coli*, and *C. acnes*. Another study found that the lower respiratory microbiome of healthy chickens is dominated by bacteria belonging to a few taxonomic classifications [17]. It was also noted that in dense living conditions and large flocks, *Avibacterium*, *Pseudomonas*, *Acinetobacter*, and *Gallibacterium* were reported as predominant genera in the lower respiratory microbiome of chickens [17].

The infected samples exhibited distinct species indicators compared to the Mock group. The samples in the SHS groups had their indicator species. Previous studies have reported that *Lactobacillus* and *Pseudomonas* genera were prevalent in the respiratory tracts of healthy chickens [17,18]. It was found that *Mycoplasma gallisepticum* (MG) infection significantly reduced the richness and diversity of respiratory microbiota compared to the control group [19]. Beta diversity analysis revealed a significant separation of microbial clusters in the buccal samples among the three groups. Therefore, considering that microbiome analysis from buccal samples is easy and noninvasive, it could be beneficial for disease screening [14,30]. Principal coordinate analysis (PCoA) indicated that samples in the MG group were markedly different from those in the control group. Meanwhile, samples from the MG and control groups were concentrated in two distinct areas [19].

The bacterial community compositions shifted after the flocks moved to the grow-out barns [20,21]. Various potential respiratory pathogens in the trachea, such as *Avibacterium*, *Erysipelothrix*, *Gallibacterium*, and *Ornithobacterium*, were found to negatively affect young broilers’ performance [22]. Additionally, the role of lung and gut microbiota in respiratory health and disease is being explored [23]. Moreover, virulence factors should be studied in the future [8,24,25]. This has implications for disease diagnosis, treatment, and vaccine development.

In summary, significant differences in microbiota composition were observed in the diseased groups (P-APG, P-ORT, P-APG-ORT, N-APG-ORT) compared to the Mock groups. Indicator species specific to the diseased groups could offer valuable insights for treating this disease. Further investigation into the unique bacteria identified in this study concerning diseased samples will contribute to developing an effective treatment against the diseases. Compared with conventional PCR, 16S rRNA gene sequencing is a promising method with high sensitivity and quantifiable for disease diagnosis. 16S rRNA gene sequencing can potentially facilitate disease control and vaccine development.

## Data available statement

Data are available from the corresponding author upon reasonable request.

## Author Contributions

Laboratory experimental design: GF Zhang. Laboratory execution of experiments, and data acquisition, analysis, and interpretation: MN Li, Q Han, N Wang, T Wang, XM You, S Zhang, CC Zhang, YQ Shi, PZ Qiao, CL Man, T Feng, YY Li, Z Zhu, KJ Quan, TL Xu, GF Zhang. Drafting and revision of the manuscript: GF Zhang. Manuscript editing and approval: GF Zhang, TL Xu, KJ Quan.

## References

[1] Y. Chen, X. Li, Y. Fu, Y. Yu, H. Zhou, Whole-genome sequencing unveils the outbreak of Mycoplasma pneumoniae in mainland China, Lancet Microbe (2024). 10.1016/S2666-5247(24)00086-7.

[2] M.A. Ali, G.F. Zhang, C. Hu, B. Yuan, S. Jahan, G.D. Kitsios, A. Morris, S.J. Gao, R. Panat, Ultrarapid and ultrasensitive detection of SARS-CoV-2 antibodies in COVID-19 patients via a 3D-printed nanomaterial-based biosensing platform, J Med Virol (2022). 10.1002/jmv.28075.

[3] M.A. Ali, G.F. Zhang, C. Hu, B. Yuan, S.J. Gao, R. Panat, An Advanced Healthcare Sensing Platform for Direct Detection of Viral Proteins in Seconds at Femtomolar Concentrations via Aerosol Jet 3D - Printed Nano and Biomaterials, Advanced Materials Interfaces (2024). 10.1002/admi.202400005.

[4] K. Quan, Z. Zhu, S. Cao, F. Zhang, C. Miao, X. Wen, X. Huang, Y. Wen, R. Wu, Q. Yan, Y. Huang, X. Ma, X. Han, Q. Zhao, Escherichia coli-Derived Outer Membrane Vesicles Deliver Galactose-1-Phosphate Uridyltransferase and Yield Partial Protection against Actinobacillus pleuropneumoniae in Mice, J Microbiol Biotechnol 28 (2018) 2095–2105. 10.4014/jmb.1809.09004.

[5] F. Zhang, S. Cao, Z. Zhu, Y. Yang, X. Wen, Y.F. Chang, X. Huang, R. Wu, Y. Wen, Q. Yan, Y. Huang, X. Ma, Q. Zhao, Immunoprotective Efficacy of Six In vivo-Induced Antigens against Actinobacillus pleuropneumoniae as Potential Vaccine Candidates in Murine Model, Front Microbiol 7 (2016) 1623. 10.3389/fmicb.2016.01623.

[6] G.F. Zhang, W. Meng, L. Chen, L. Ding, S. Sun, X. Wang, Y. Huang, H. Guo, S.J. Gao, Infectivity of pseudotyped SARS-CoV-2 variants of concern in different human cell types and inhibitory effects of recombinant spike protein and entry-related cellular factors, J Med Virol 95 (2023) e28437. 10.1002/jmv.28437.

[7] F. Zhang, Q. Zhao, K. Quan, Z. Zhu, Y. Yang, X. Wen, Y.F. Chang, X. Huang, R. Wu, Y. Wen, Q. Yan, Y. Huang, X. Ma, X. Han, S. Cao, Galactose-1-phosphate uridyltransferase (GalT), an in vivo-induced antigen of Actinobacillus pleuropneumoniae serovar 5b strain L20, provided immunoprotection against serovar 1 strain MS71, PLoS One 13 (2018) e0198207. 10.1371/journal.pone.0198207.

[8] F. Zhang, Y. Zhang, X. Wen, X. Huang, Y. Wen, R. Wu, Q. Yan, Y. Huang, X. Ma, Q. Zhao, S. Cao, Identification of Actinobacillus pleuropneumoniae Genes Preferentially Expressed During Infection Using In Vivo-Induced Antigen Technology (IVIAT), J Microbiol Biotechnol 25 (2015) 1606–1613. 10.4014/jmb.1504.04007.

[9] M. Morsli, Y. Bechah, O. Coulibaly, A. Toro, P.E. Fournier, L. Houhamdi, M. Drancourt, Direct diagnosis of Pasteurella multocida meningitis using next-generation sequencing, Lancet Microbe 3 (2022) e6. 10.1016/S2666-5247(21)00277-9.

[10] J.B. Patel, 16S rRNA gene sequencing for bacterial pathogen identification in the clinical laboratory, Mol Diagn 6 (2001) 313–321. 10.1054/modi.2001.29158.

[11] P.C. van Empel, H.M. Hafez, Ornithobacterium rhinotracheale: A review, Avian Pathol 28 (1999) 217–227. 10.1080/03079459994704.

[12] H.M. Hafez, Diagnosis of Ornithobacterium Rhinotracheale, International Journal of Poultry Science 1 (2002) 114–118. 10.3923/ijps.2002.114.118.

[13] M.N. Li, T. Wang, N. Wang, Q. Han, X.M. You, S. Zhang, C.C. Zhang, Y.Q. Shi, P.Z. Qiao, C.L. Man, T. Feng, Y.Y. Li, Z. Zhu, K.J. Quan, T.L. Xu, G.F. Zhang, A detailed analysis of 16S rRNA gene sequencing and conventional PCR-based testing for the diagnosis of bacterial pathogens and discovery of novel bacteria, Antonie Van Leeuwenhoek 117 (2024) 102. 10.1007/s10482-024-01999-1.

[14] B. Goldberg, H. Sichtig, C. Geyer, N. Ledeboer, G.M. Weinstock, Making the Leap from Research Laboratory to Clinic: Challenges and Opportunities for Next-Generation Sequencing in Infectious Disease Diagnostics, mBio 6 (2015) e01888–01815. 10.1128/mBio.01888-15.

[15] M. Fida, S. Khalil, O. Abu Saleh, D.W. Challener, M.R. Sohail, J.N. Yang, B.S. Pritt, A.N. Schuetz, R. Patel, Diagnostic Value of 16S Ribosomal RNA Gene Polymerase Chain Reaction/Sanger Sequencing in Clinical Practice, Clin Infect Dis 73 (2021) 961–968. 10.1093/cid/ciab167.

[16] I. Rychlik, D. Karasova, M. Crhanova, Microbiota of Chickens and Their Environment in Commercial Production, AVIAN DISEASES 67 (2023) 1–9.

[17] M.Z. Shabbir, T. Malys, Y.V. Ivanov, J. Park, M.A. Shabbir, M. Rabbani, T. Yaqub, E.T. Harvill, Microbial communities present in the lower respiratory tract of clinically healthy birds in Pakistan, Poult Sci 94 (2015) 612–620. 10.3382/ps/pev010.

[18] P.E. Larsen, L. Glendinning, G. McLachlan, L. Vervelde, Age-related differences in the respiratory microbiota of chickens, Plos One 12 (2017). 10.1371/journal.pone.0188455.

[19] J. Wang, M. Ishfaq, Q. Fan, C. Chen, J. Li, A respiratory commensal bacterium acts as a risk factor for Mycoplasma gallisepticum infection in chickens, Vet Immunol Immunopathol 230 (2020) 110127. 10.1016/j.vetimm.2020.110127.

[20] K.J.M. Taylor, J.M. Ngunjiri, M.C. Abundo, H. Jang, M. Elaish, A. Ghorbani, M. Kc, B.P. Weber, T.J. Johnson, C.W. Lee, Respiratory and Gut Microbiota in Commercial Turkey Flocks with Disparate Weight Gain Trajectories Display Differential Compositional Dynamics, Appl Environ Microbiol 86 (2020). 10.1128/AEM.00431-20.

[21] J.M. Ngunjiri, K.J.M. Taylor, M.C. Abundo, H. Jang, M. Elaish, M. Kc, A. Ghorbani, S. Wijeratne, B.P. Weber, T.J. Johnson, C.W. Lee, Farm Stage, Bird Age, and Body Site Dominantly Affect the Quantity, Taxonomic Composition, and Dynamics of Respiratory and Gut Microbiota of Commercial Layer Chickens, Appl Environ Microbiol 85 (2019). 10.1128/AEM.03137-18.

[22] J.M. Diaz Carrasco, N.A. Casanova, M.E. Fernandez Miyakawa, Microbiota, Gut Health and Chicken Productivity: What Is the Connection?, Microorganisms 7 (2019). 10.3390/microorganisms7100374.

[23] T.P. Wypych, L.C. Wickramasinghe, B.J. Marsland, The influence of the microbiome on respiratory health, Nature Immunology 20 (2019) 1279–1290. 10.1038/s41590-019-0451-9.

[24] Z. Zhu, Q. Zhao, Y. Zhao, F. Zhang, X. Wen, X. Huang, Y. Wen, R. Wu, Q. Yan, Y. Huang, X. Ma, X. Han, S. Cao, Polyamine-binding protein PotD2 is required for stress tolerance and virulence in Actinobacillus pleuropneumoniae, Antonie Van Leeuwenhoek 110 (2017) 1647–1657. 10.1007/s10482-017-0914-7.

[25] F. Zhang, Q. Zhao, J. Tian, Y.F. Chang, X. Wen, X. Huang, R. Wu, Y. Wen, Q. Yan, Y. Huang, X. Ma, X. Han, C. Miao, S. Cao, Effective Pro-Inflammatory Induced Activity of GALT, a Conserved Antigen in A. Pleuropneumoniae, Improves the Cytokines Secretion of Macrophage via p38, ERK1/2 and JNK MAPKs Signal Pathway, Front Cell Infect Microbiol 8 (2018) 337. 10.3389/fcimb.2018.00337.

